# *k*-mer grammar uncovers maize regulatory architecture

**DOI:** 10.1101/222927

**Authors:** María Katherine Mejía-Guerra, Edward S Buckler

**Affiliations:** Institute for Genomic Diversity, Cornell University, Ithaca, NY 14853; USDA-ARS, Research Geneticist, USDA ARS Robert Holley Center, Ithaca, NY 14853; Department of Plant Breeding and Genetics, Cornell University, Ithaca, NY 14853

**Author notes:** Correspondence: María Katherine Mejía-Guerra.

## Abstract

Only a small percentage of the genome sequence is involved in regulation of gene expression, but to biochemically identify this portion is expensive and laborious. In species like maize, with diverse intergenic regions and lots of repetitive elements, this is an especially challenging problem. While regulatory regions are rare, they do have characteristic chromatin contexts and sequence organization (the grammar) with which they can be identified. We developed a computational framework to exploit this sequence arrangement. The models learn to classify regulatory regions based on sequence features - *k*-mers. To do this, we borrowed two approaches from the field of natural language processing: (1) “bag-of-words” which is commonly used for differentially weighting key words in tasks like sentiment analyses, and (2) a vector-space model using word2vec (vector-*k*-mers), that captures semantic and linguistic relationships between words. We built “bag-of-*k*-mers” and “vector-*k*-mers” models that distinguish between regulatory and non-regulatory regions with an accuracy above 90%. Our “bag-of-*k*-mers” achieved higher overall accuracy, while the “vector-*k*-mers” models were more useful in highlighting key groups of sequences within the regulatory regions. These models now provide powerful tools to annotate regulatory regions in other maize lines beyond the reference, at low cost and with high accuracy.

## INTRODUCTION

The vast majority of sequence polymorphisms that are statistically associated with phenotypic variation (GWAS) lie in the non-genic portion of the genome, where they might play regulatory roles [1,2]. Recently biochemical characterization of the open chromatin space in B73 (the maize reference line), revealed that as much as 40% of the significant sequence polymorphisms - as identified through variance components analyses – overlap with regions in which regulatory elements are expected [3]. These biochemical assays are prohibitively expensive and time consuming at the scale of breeding programs for any crop species. This is even more true for species, such as maize, with high genomic diversity and a high rate of polymorphism. In maize, less than half of the genome sequence is expected to be shared between inbred lines [4]. Building accurate models from expensive data derived from the maize reference line will enable breeders to broadcast that information to other genotypes for use in genomic selection models and to prioritize regions of the genome to edit using strategies such as CRISPR [5,6].

Most of the experimental and computational approaches used to annotate functional noncoding regions focus on the regulatory role of transcription factor binding sites (TFBSs) [7,8]. However, it has been observed that patterns of sequence organization (the grammar) and the chromatin context in which TFBSs are located contribute to the regulatory message[9–11]. For instance, the spatial arrangement of poly(dA:dT) tracts within yeast promoter regions have been identified as causal drivers of transcriptional patterns at comparable levels to TFBSs [12]. More recently, it was shown that developmental enhancers in *Ciona* rely on the positioning, arrangement, and space between TFBSs to counterbalance low TFBS affinity [13]. From this emerging view, it appears that regulatory regions have distinctive features that can be exploited for prediction, identifying enriched key sequences and sequence organization.

The frequency of oligomers of length *k (i.e*., short *k*-mers in the size range of TFBS) have been exploited to build supervised models capable of discriminating regulatory regions from random genomic regions, as well as to score sequence variation with few or no assumptions regarding to the role that a given *k*-mer might play [14–16]. The early *k*-mer count-based classifiers have been improved to count gapped *k*-mers, allowing exploration of short and long *k* values without losing power as the total number of *k*-mers increases [17]. Some limitations of *k*-mer frequency-based methods include: (1) they make poor or no use of the *k*-mer positional relationships in their models, and (2) they perform poorly in the presence of repetitive regions, the frequencies of short size *k*-mer are misleading, which might hamper the performance of this methods for genomes with high repeat content.

Recently however, a growing set of computational tools using Neural Networks (NNs) have shown success in learning to recognize simple sequence patterns, similar to Position Weight Matrices (PWMs). These approaches have been able to further integrate those patterns into more complex features to discriminate regulatory regions [18–20]. Generally, the NNs implemented for genomic data are Convolutional Neural Networks (CNNs), a type of architecture that shows state-of-the-art performance for key phrase recognition tasks in Natural Language Processing (NLP), but not Recurrent Neural Networks (RNNs) which are preferred for comprehension of whole sentence semantics given their power in modeling long-span relations [21–22]. Despite their power, CNNs are often implemented in a black-box context and interpretation of their output is challenging; thus it remains unclear how much of their performance is derived from recognizing key motifs, motif relationships, and the general sequence context. For these reasons we choose to implement *k*-mer approaches rather than CNN’s or RNN’s.

To define groups of *k*-mers sharing regulatory roles, we analyzed the architecture of regulatory regions at the *k*-mer level, focusing on weighted individual frequencies and cooccurrences. The core of the analysis builds on NLP machine learning approaches that are easily interpretable and rely on word statistics to recover semantic and syntactic cues [23–26]. We evaluated the accuracy and precision of these approaches with a diverse set of functional genomics experiments to provide a comprehensive description of the regulatory landscape of the maize genome. The software implementation is open source and available through Bitbucket repository.

## RESULTS

### Weighted frequencies and co-occurrences of short sequences can accurately discriminate regulatory from random genomic regions

To build accurate classifiers we collected a comprehensive set of regions enriched in regulatory function (hereafter, ‘regulatory regions’), as identified in B73 (maize reference genome) through different biochemical assays. We included in the analysis open chromatin regions by MNAseq derived from two tissues [3], binding loci from ChIP-seq peaks of two TFs (*i.e*., Homeobox Knotted 1 – KN1, bZIP Fasciated ear4 – FEA4) [27,28], and core promoter regions around TSSs [29–31] (Supplementary Table 1). Because the specific background signals from each individual experiment are not available, regulatory regions were paired with randomly chosen regions controlling for G+C content and genomic distribution. Each group of sequence (regulatory regions and their control) was separated into training and holdout sets for model evaluation. In total we analyzed 52,292,705 bp of regulatory regions corresponding to ~2.5% of the B73 genome.

The first part of the analysis involved the training of “bag-of-*k*-mers” and “vector-*k*-mers” models (Figure 1). The “bag-of-*k*-mers” captures information from the *k*-mer individual frequencies and fits a logistic regression to a matrix filled with the TF*IDF transformation of the frequencies per sequence [23]. Thus, the β coefficients of the logistic regression can be interpreted as weights of the contribution of each *k*-mer to the classifier decision and of its enrichment in regulatory and random regions. By contrast, the “vector-*k*-mers” captures information from the *k*-mer co-occurrences by training a shallow NN that learns the probability for each *k*-mer given its context (window = *5*). The output is 300 dimensional vectors (*v*_k-mer_) – one per *k*-mer - independently generated for regulatory regions and their respective control (*V*_regulatory_ and *V*_random_ to denote different geometric spaces containing *v_k-mers_*). Next, *V*_regulatory_ and *V*_random_ are utilized to determine the likelihood of groups of *k*-mers being observed in regulatory or control regions [25,26]. Put together, these two models aim to learn the importance of key sequence features and sequence feature relationships as descriptors of regulatory architecture.

**Figure 1:**
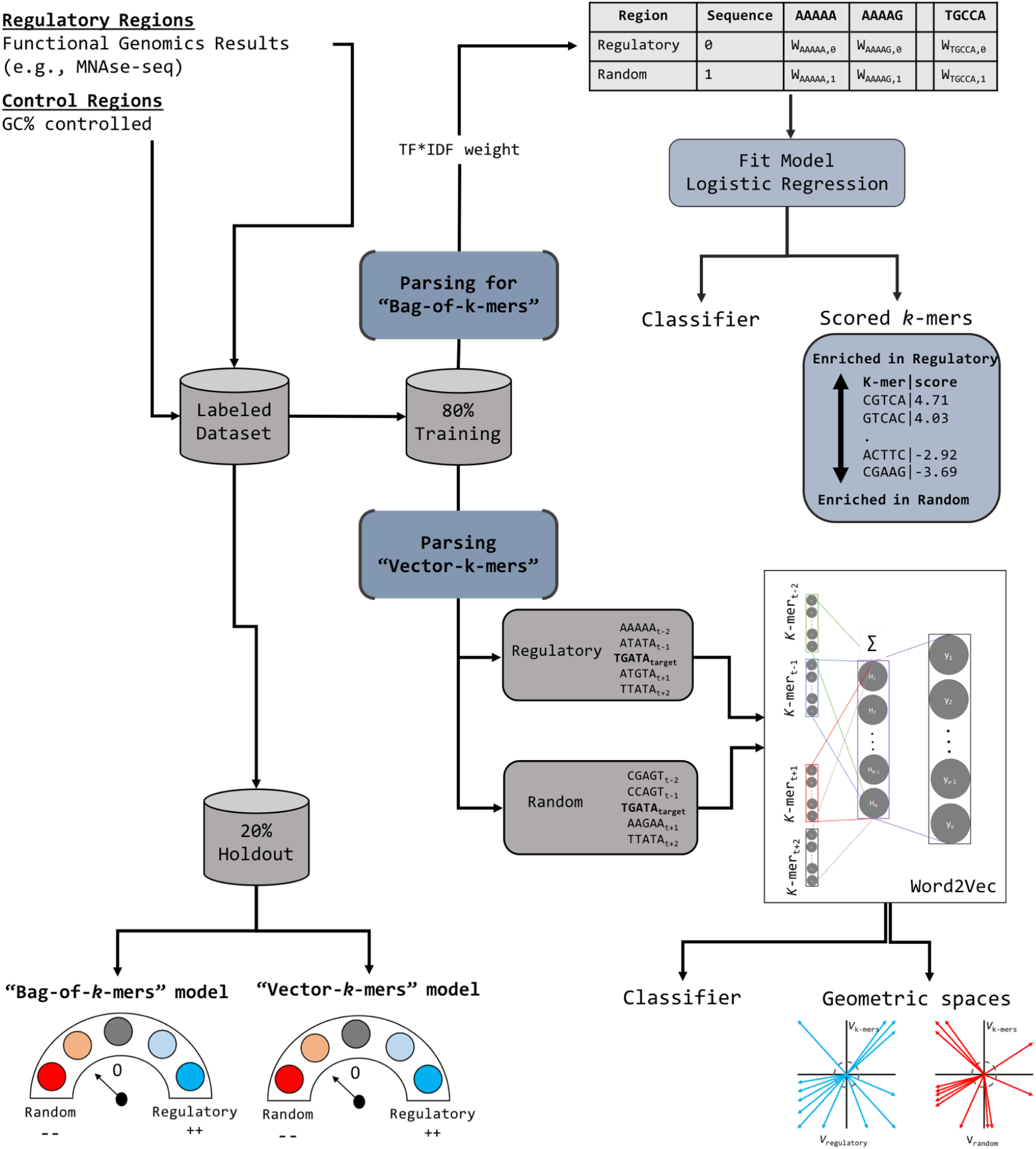
Schematic of the Steps to Generate “bag-of-*k*-mers” and ‘Vector-*k*-mers” Models. The above workflow shows the steps from data preprocessing tomodel output. We fitted “bag-of-*k*-mers” and “vector-*k*-mers” models for *k* values between 5 to 10 bp (within the common range in which regulatory elements have been observed). Training and evaluaticn of both methods happened on the same portion of the data to facilitate comparisons. The common pre-processing step involved the collapsing of complementary *k*-mers as the same token to reduce the noise of *k*-mer counts and the effective vocabulary for feature selection. The final outputs are both the classifiers and learned features.

Model performance was measured with three metrics: (1) accuracy, (2) the area under the receiver operating characteristic (auROC) curve, and (3) the area under the precision recall curve (auPRC) (Supplementary Table 2). The first evaluation was done on balanced holdout sets (*i.e*., the same number of regulatory and random sequences) (Fig. 2a-d). The two models perform similarly, with an average difference of 3% in accuracy between the two for any given *k*. Overall, the “bag-of-*k*-mers” model shows better performance for most of the cases, with the “vector-*k*-mers” models only outperforming when *k* is small (*k*=5 and *k*=6) and training datasets are large (*e.g*., MNAseq shoot) (Supplementary Table 3). The performance of the “bag-of-*k*-mers” models was reliable even at *k* ≥ 8, as opposed to similar approaches that rely on raw *k*-mer counts as features to train machine learning classifiers [15,32]. The above suggests that the TF*IDF transformation is efficient in alleviating some of the noise inherent to the matrix sparsity that increased with *k*.

**Figure 2:**
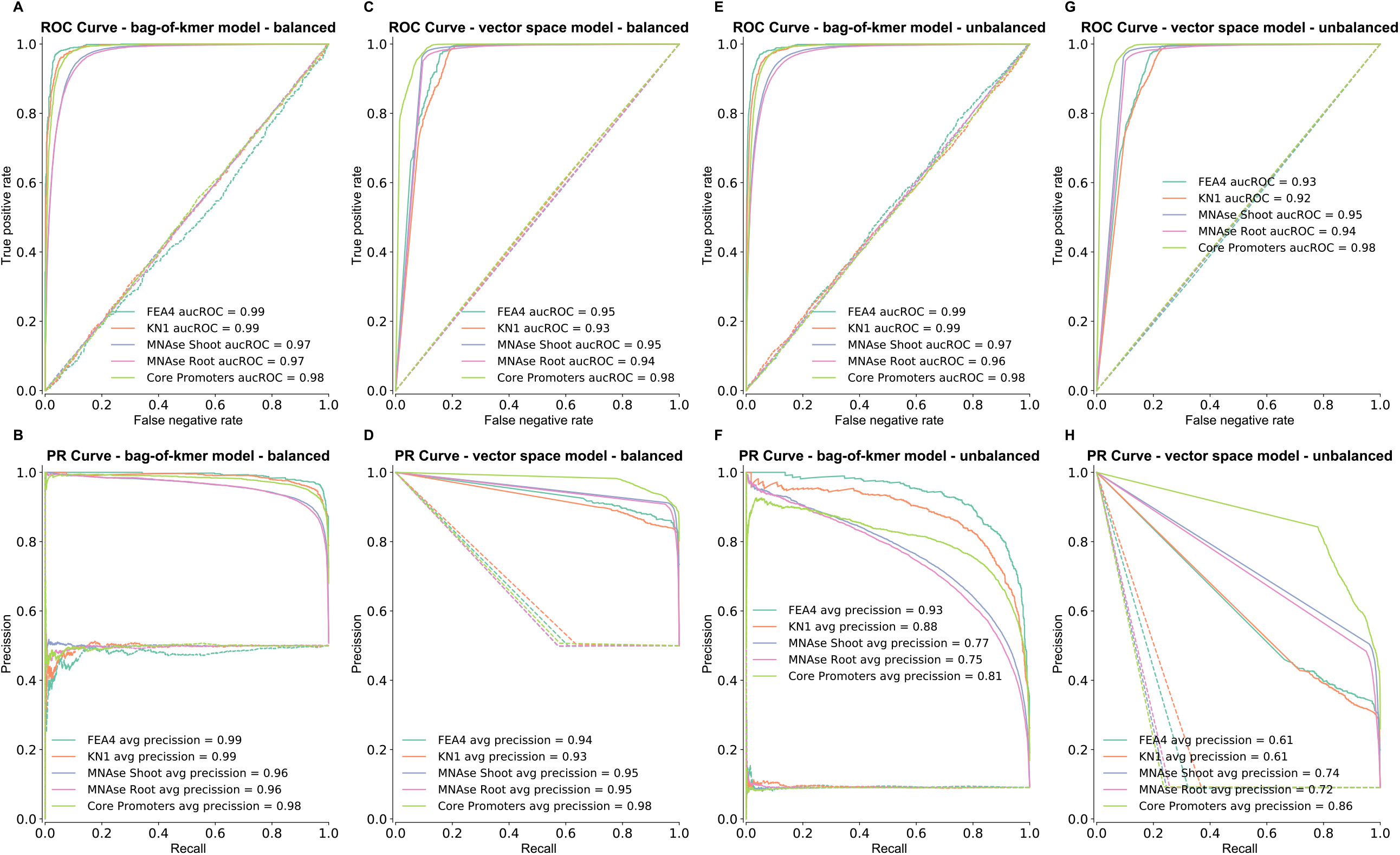
Comparison ofROC and PRC Curves for Prediction of Regulatory Regions. Comparison of models performance under balanced (**a - d**) and unbalanced holdout sets (**e - h**). For each model (*k*=8), the receiver operating characteristic (ROC) curve and the precision recall curve (PRC) are shown for all the regulatory datasets and the corresponding curves for classification of the holdout set with randomized labels (dotted lines)

To increase the stringency of our evaluation criteria, we measured each models’ performance with unbalanced holdout sets in which regulatory regions are outnumbered by random regions by 1 to 10 (Fig. 2e-h, Supplementary Table 4). Scaling up the number of random regions did not appreciably change accuracy and auROC values, but the auPRC showed a drop in model performance as the rate of false positive increased. At *k*=8, both models have a desirable precision, ~80-70%, at a desirable recall rate of ~60% for open chromatin and core promoter datasets. The “bag-of-*k*-mers” model works better for prediction of TF binding loci than the “vector-*k*-mers”, with the last one displaying an excess of false positives at our aimed recall rate. The performance measurement under an unbalanced set suggests that applying extra stringency to the predicted probability, thereby allowing the recovery of ~60% of the relevant sequences, would result in an acceptable tradeoff between sensitivity and specificity for most of the models when non-regulatory regions are in large numbers.

Highly repetitive genomes include an abundance of low-complexity regions. These repetitive regions are expected to carry little information for regulation, and because of their high-frequency, they represent an obstacle to identifying the key elements from raw *k*-mer counts. To empirically determine a complexity threshold for *k*-mers unlikely to have a regulatory role, we examined a collection of regulatory motifs and calculated complexity (as measured with Shannon entropy) for the consensus sequences (Supplementary Fig. 1). Using this threshold, *k*-mers with low complexity were filtered out to build “bag-of-*k*-mers” models with a reduced vocabulary (filtered), and contrasted against models using the whole vocabulary (full). The differences between the two models at a base pair level is illustrated for the *ga2ox1* first intron recognized by KN1 [27,33]. We observed that low complexity regions overlapped with *k*-mers that have a high score from the model trained on the full *k*-mer vocabulary (Fig. 3a). This is different from the filtered model which appears to be in agreement with the ChIP-seq data (Fig. 3, Supplementary Fig. 2). To evaluate the importance of these repetitive sequences in recognizing the regulatory regions, we compared the models with and without low complexity *k*-mers using an unbalanced holdout set and found that both models show almost identical performance, as observed from the comparison of the auROC (Fig. 2c, 3b) and the auPRC curve (Fig. 2g, 3c). This suggests that in general, low complexity *k*-mers in maize do not contribute substantially to the regulatory message.

**Figure 3:**
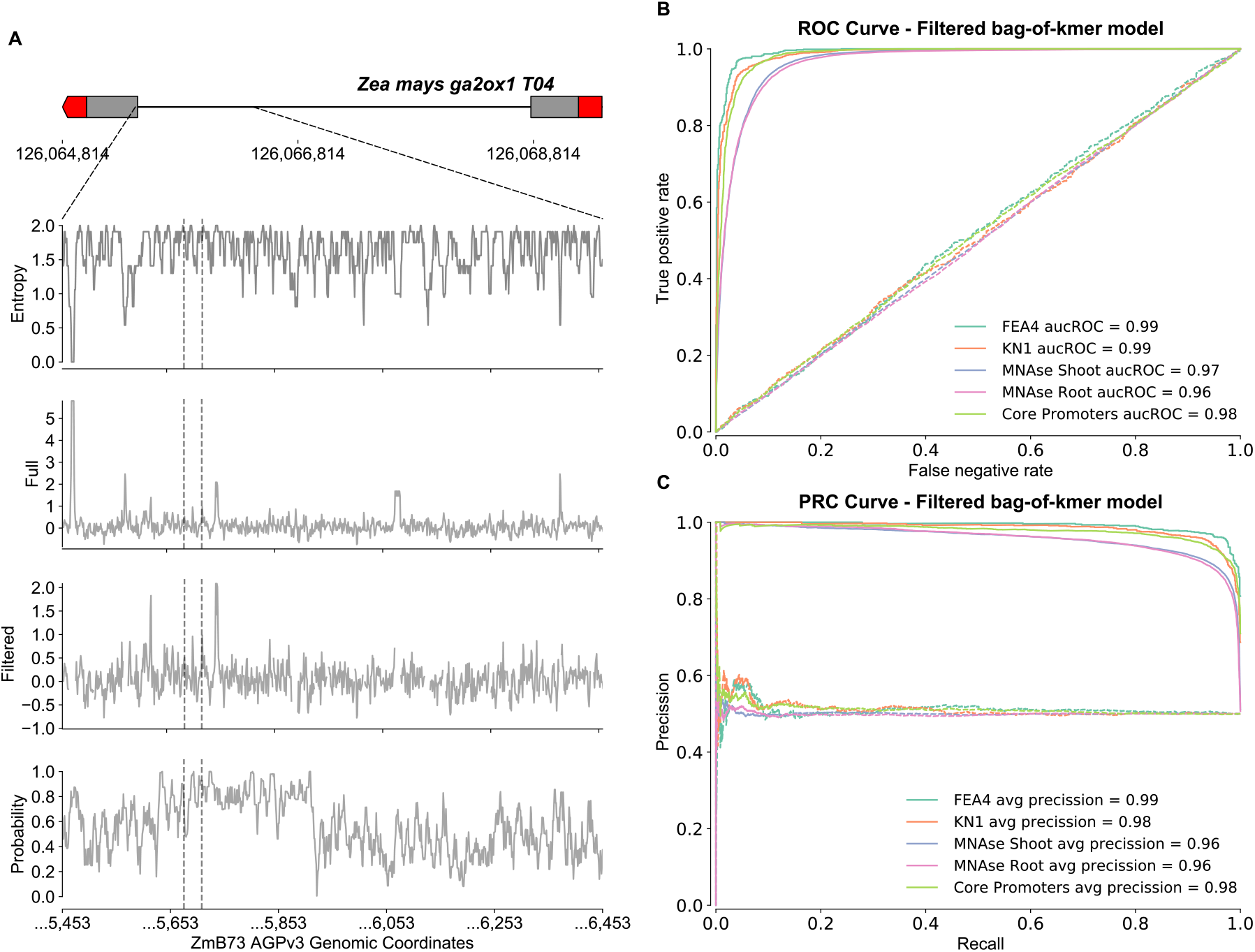
Low Complexity Regions do not Provide Relevant Information to Discriminate Regulatory Regions. (**a**) Annotation at a base pair level of the first 1Kb of the long intron in the maize gene *ga2ox1* using sequence complexity (Entropy), scores from “bag-of-*k*-mers” models (Full and Filtered), and regulatory probabilities (Probability) from the “vector-*k*-mers” model. Sequence complexity and “bag-of-*k*-mers” scores were calculated using a 1bp sliding window of size *k*. Regulatory probabilities were calculated using a 1bp sliding window of 3\**k* to evaluate co-occurrence of groups of 3 and 2 *k*-mers. The evaluated region includes the KN1 ChIP-seq peaks as identified from two biological replicates in developing ears (the center of the peak for each replicate is indicated with a vertical dotted line). (**b**) ROC curve and (**c**) PRC curve are shown for a “bag-of-*k*-mers” model (*k*=8) after removal of low-complexity *k*-mers (filtered) and tested with unbalanced holdout set.

### Models to predict regulatory regions are scalable to the genome-wide space and transferable to related species

Under the assumption that annotation of non-coding regions would be part of general pipelines, in which ~85% of the genome should be recognized as repeats and ~5% as coding sequences, our models for annotating regulatory regions should be limited to ~10% of the space. To adhere to the above scenario while gaining insights on the behavior of the models at a genome-wide scale, the sequence of chromosome 10 was partitioned into 1,943,698 regions (300 bp length) and 115,149 regions that were neither repeats nor coding sequences were selected to be annotated. We used models derived from MNAseq shoot data applying different levels of stringency for the predicted probabilities (Supplementary Table 4). According to the results obtained with unbalanced holdout set, and in order to balance sensitivity and specificity, we determined that the ideal predicted probability cut-off was the one that captures ~60% of the regions that overlap with the annotated regulatory regions. Under this criteria the “bag-of-*k*-mers” (*k*=8, filtered, probability > 0.85) and the “vector-*k*-mers” models (probability > 0.95), predicted 38,945 and 41,932 regulatory regions respectively. The high confidence regions classified as regulatory correspond to ~2.2-2.3% of the total regions from chromosome 10, in line with the expected portion of the genome with a regulatory function.

Transference of functional genomic annotations across diverse maize lines requires models than can preferentially capture conserved features (those common between lines or related species). Consistently, we expect that models that are accurate in related species should also perform well in different maize lines. To gain insights into this we evaluated models trained on core promoters and TF binding loci in two species (sorghum and rice). For the evaluation of models trained on core promoters we used a balanced holdout set derived from a random sample of sorghum annotated gene models. We obtained auROC values of 0.72 and 0.63 for the “bag-of-*k*-mers” (*k*=8, filtered) and “vector-*k*-mers” models, respectively, which represent a reduction of ~30% compared to the maize holdout data, as a result of a higher false positive rate (Fig. 4a-d). This might be a consequence of the strong differences between the repeat landscape in the non-coding regions between sorghum and maize that is not captured in the maize training set, rather than a lack of similarities between the regulatory regions of the two species.

**Figure 4:**
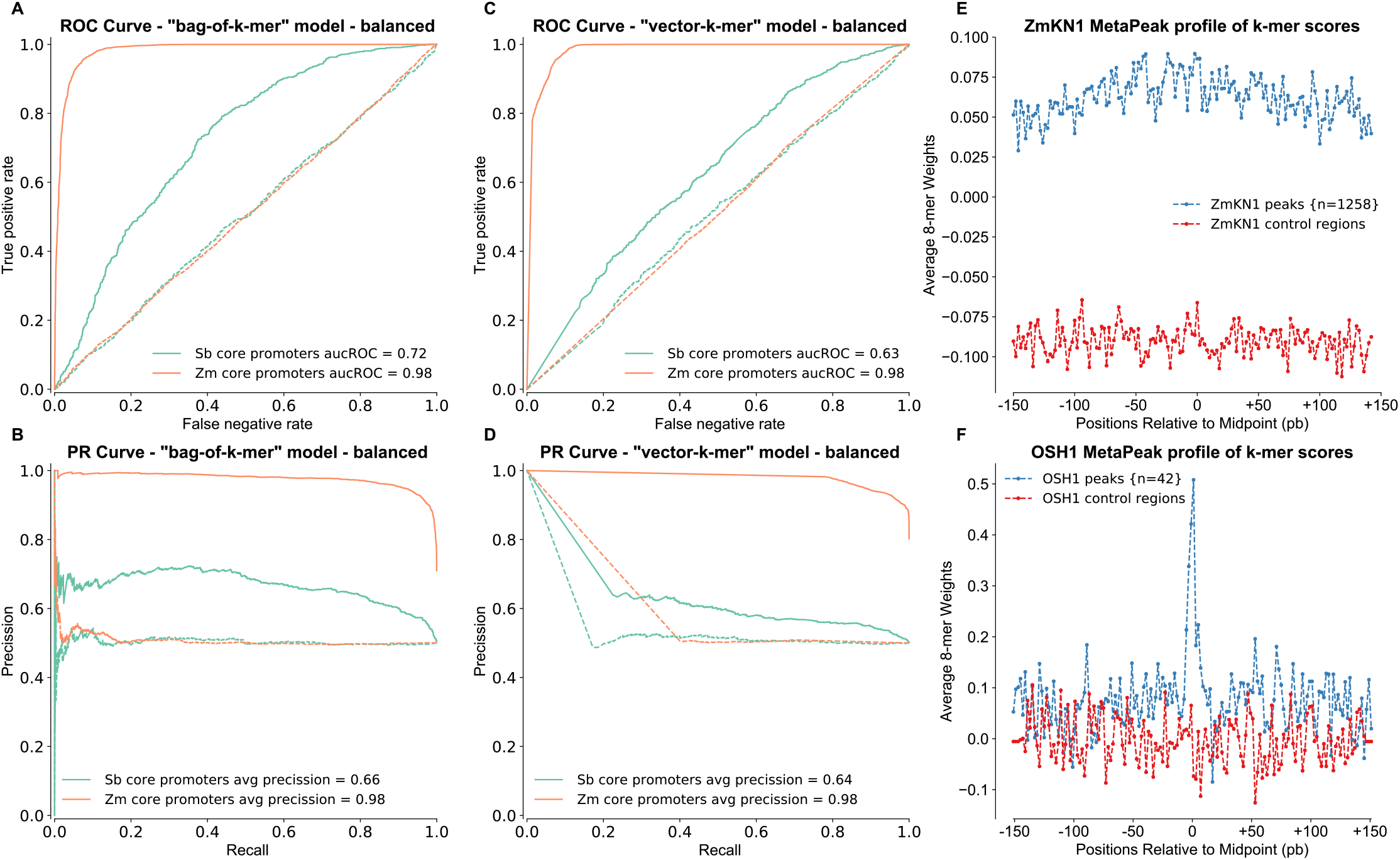
“bag-of-*k*-mers” and “vector-*k*-mers” Models can Accurately Predict Core Promoter Regions and TF Binding Loci Across Species. ROC curve and PRC curves summarizing the performance of “bag-of-*k*-mers” (filtered) (**a-b**) and “vector-*k*-mers” (**c-d**) models (*k*=8) trained in maize core promoter and tested in a set of randomly chosen 1,000 core promoter regions and their respective controls in sorghum. (**e**) Base-pair annotation of KN1 binding loci (blue) and control regions (red) using scores derived from a “bag-of-*k*-mer” model (filtered) trained on KN1 (Knotted 1, ZmHD1) ChIP-seq data [27] show a slight preference for the center of the peak over flanking regions. (**f**) The same model also works in rice, differentiating regions targeted by OSH1 (the functional orthologue of KN1) [34] (blue) from control regions (red) with the same GC content with a strong preference for the center of the peak.

In order to determine positional preferences among binding loci, we built peak metaprofiles that summarized KN1 models’ performance in maize and rice at the base-pair level (Fig. 4e-f, Supplementary Fig. 3a-b). The “bag-of-*k*-mers” model can differentiate between regulatory regions and their control in maize, and in addition can distinguish rice KN1-like (*i.e*., OSH1) binding sites (*i.e*., peaks from rice OSH1 ChIP-seq data [34]). On the other hand, the “vector-*k*-mers” cannot differentiate between random regions and regulatory regions in rice, predicting random as regulatory (Supplementary Fig. 3a-b). Interestingly, the distributions of regulatory probabilities for random and regulatory regions are noticeable different (Supplementary Fig. 3c), suggesting that the “vector-*k*-mers” model distinguish between OSH1 peaks and control regions, but not enough to assign greater non-regulatory probability to random regions. In maize, the “bag-of-*k*-mers” model (filtered) shows a slight preference towards the midpoint region versus the edges, while the “vector-*k*-mers” recognizes the whole region without preference for to the middle (Fig. 4e). In rice, the “bag-of-*k*-mers” shows a marked preference near or at the peak midpoint over the flanking (Fig. 4f). This suggests that “bag-of-k-mers” models capture a diverse array of features which are enriched at the center of the peak and beyond in maize. However, only the key features that are enriched at the center of the peak appear indeed conserved between the two species. Taken together we have shown that classifiers trained in maize can be useful to predict regulatory regions in sorghum and rice, and that features enriched in maize regulatory regions and in the random genomic space (as captured by the models) are of two general types: (1) maize specific and (2) conserved across related species.

### Scored vocabularies highlight signatures of regulatory function

The methods proposed here were chosen because of the interpretability of the learned features, aiming to better understand the patterns in sequence that characterize regulatory regions. Thus, we focused on scored *k*-mer vocabularies (*k*=8, filtered) as easiest to interpret, and systematically analyzed the tails of the distribution as they concentrated the most informative sequences. Therefore, the largest positive coefficient values (top scored *k*-mers) are indicative of enrichment and the largest negative values (bottom scored *k*-mers) of depletion in regulatory regions. The absolute values from both sides of the score distribution are different, with preference for positive over negative ones, meaning that model’s prediction are the result of identifying those *k*-mers that are enriched in regulatory regions rather than depleted ones (or enriched in random regions). We found that properties of the scored *k*-mers obtained from applying an out-of-the-box NLP technique [23] are similar to those previously described with sequence kernels developed to analyze vertebrate genomic data [15,17].

We observed a bias in the G+C content at the extremes of the score distribution for core promoters (Fig. 5a) and to a lesser extend for open chromatin regions (Fig. 5b-c). The 1% of the top shows a bimodal distribution, in which a subpopulation of *k*-mers exhibits low G+C content, in contrast to the 1% of the bottom, and the remaining 98%. Conversely, the score distribution for TF binding loci shows a general shift of top and bottom tails towards higher G+C contents, in comparison to the remaining 98% (Supplementary Fig. 4). These results are in agreement with known roles for high A+T sequences within core promoters related to the TATA elements and high G+C sequences as TF binding sites [31,35]. Indeed, when investigated, individual *k*-mers with high A+T content were positionally restricted upstream of the TSS and preferentially on the region defined for the TATA element in maize (Fig. 5d).

**Figure 5:**
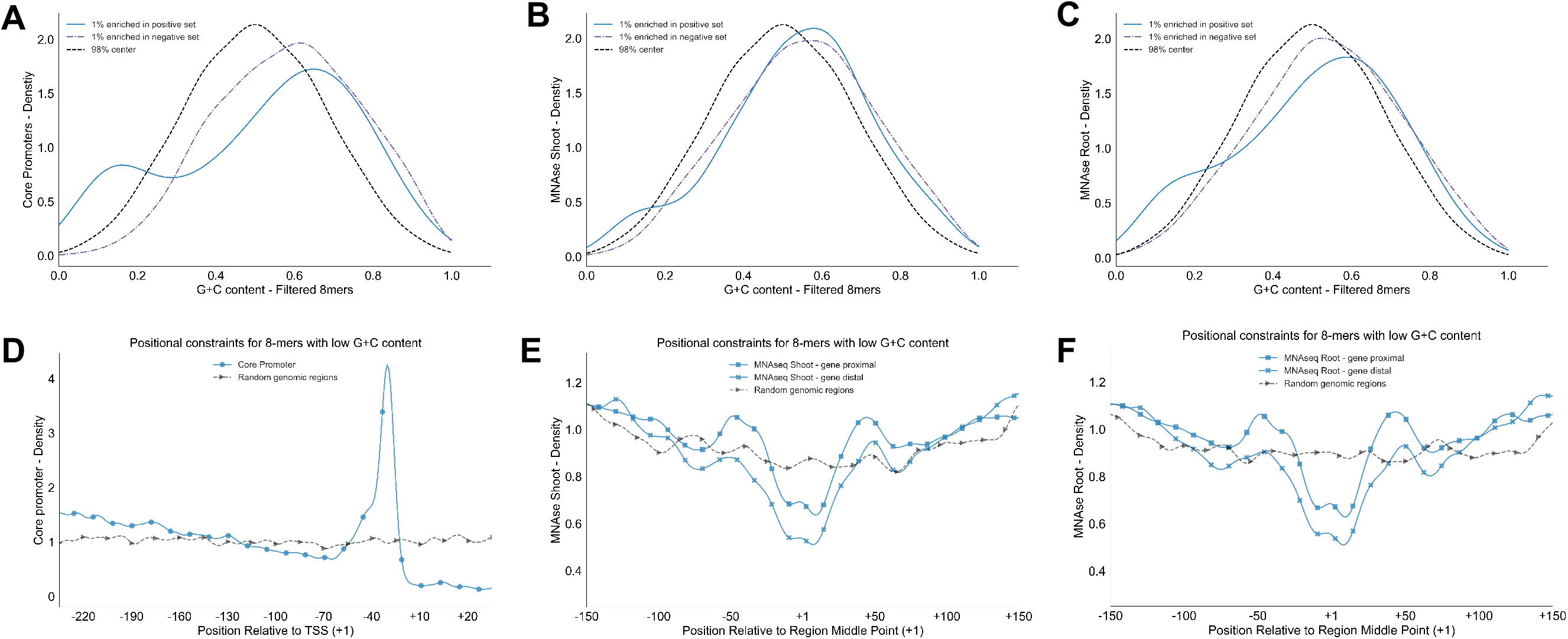
Scored Vocabularies Uncovers G+C Content Bias Positionally Constrained within Promoters and Enhancer Regions. Comparison of the distribution of G+C content across top 1%, bottom 1% and remaining 98% of scored *k*-mer vocabularies (*k*=8, filtered) for (**a**) core promoters, MNAseq (**b**) shoot and (**c**) root model’s results. The positional constraints of *k*-mers with high A+T content on the top 1% visualized as *k*-mer’s density with respect to a reference point: (**d**) TSS for core promoters and MNAseq hotspot middle point (**e**) shoot and (**f**) root (solid blue lines). Contrasting density plots are shown for corresponding random regions (dotted gray lines).

The enrichment of MNAseq regions for *k*-mers with high A+T content (rich A+T *k*-mers) might be derived from signal co-localization between open chromatin regions and core promoters [3]. If signal co-localization were sufficient to explain the similarities between open chromatin and core promoter regions, then controlling for distance to annotated genes should remove the signal from rich A+T *k*-mers in distal regions. However, even when gene proximal regions account for more signal than distal ones, the positional constraints remain in both proximal and distal regions (Fig. 5e-f). These rich A+T *k*-mers might be part of poly(dA:dT) tracts which can provide an increase in DNA rigidity and are known to be in proximity to regions that are enriched in TFBSs [36]. In agreement with the positional restriction, rich A+T *k*-mers flank the midpoints where G+C content is high, as expected for the regions that are bound by TFs [35], and where the signal for open chromatin regions is concentrated.

In addition to key structural tracts, *k*-mers with the largest positive values for each regulatory category are expected to be enriched for TF motifs. Because the number of experimentally verified maize motifs is limited, we contrasted the top 1% of positive scored *k*-mers against two large collections of TF motifs as identified from large scale experiments in the reference plant *Arabidopsis thaliana* (TOMTOM, *p*-value < 0.001 [39]) [38–39]. For the evaluated experiments we found that the top 1% of positive *k*-mers are ~threefold more enriched for significant hits against the motif database than expected by chance for all the *k*-mers in the population. The enrichment for the top *k*-mers was statistically significant (hypergeometric test, *p*-value < 0.001). Further analyses revealed that *k*-mer scoring is consistent within families of TF binding sites. In particular, motifs preferentially hit by the top 1% of positive *k*-mers from FEA4 binding loci (a bZIP transcription factor) correspond to the bZIP/TGA-class, and motifs preferentially hit by *k*-mers enriched in KN1 (a Homeobox transcription factor) correspond to the Homeobox family (Supplementary Table 5). Thus, the scored vocabularies produced a comprehensive catalog of *k*-mers with putative structural roles and a collection of *k*-mers similar to TFBSs that constitute signatures of the maize regulatory architecture.

### Sequence similarity in the geometric space reveals a prevalent distinctive *k*-mer organization within regulatory regions

The set of highly enriched individually scored sequences, as output from “bag-of-*k*-mers” models, is likely to include groups of *k*-mers that correspond to the same motif, given the degeneracy of TFs binding sites. However, the question arises of how to group *k*-mers that likely share functional roles and constitute single motifs. In NLP, Word2Vec is an effective method to extract linguistic regularities between words by considering the local context in which they occurs (*e.g*., apple and oranges might share local contexts as they are words with similar meanings) [40]. Because vector position in each geometric space is determined from the composition of the local word/*k*-mer context (*i.e*., neighboring *k*-mers), we can assume that two *k*-mers that are close (*i.e*., close in cosine distance) to each other in a geometric space share local sequence similarity (Fig. 6a). Therefore, we used the geometric spaces obtained from the “vector-*k*-mers” models, to extract *k*-mer regularities or *k*-mer organizational ‘rules’ that differentially arise between regulatory and random regions. Because, the position of *k*-mers between geometric spaces cannot be directly contrasted, we compared the lists of closest *k*-mers for any given *k*-mer in the vocabulary as obtained from the geometric spaces about regulatory and random regions (respectively, *V_reguiatory_* and *V_random_*).

**Figure 6:**
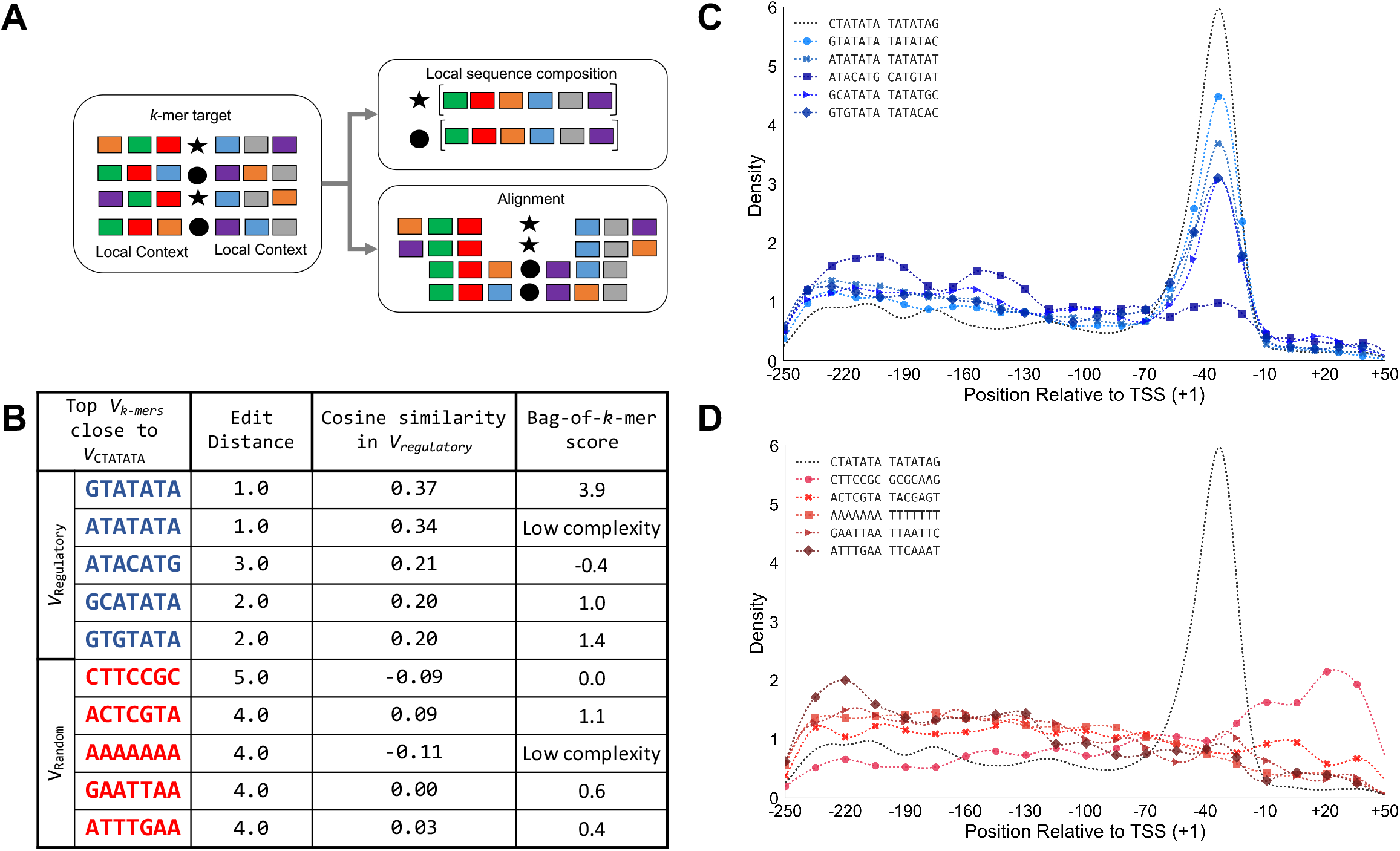
“vector-*k*-mers” Models Capture Related *k*-mers from the Influence of their Local Sequence Context Within Regulatory Regions. (**a**) Schematic of capturing local sequence composition versus aligning flanking contexts as implemented in the “vector-*k*-mers” models in which *k*-mers that share a similar context would be represented by close vectors (*v_k-mers_*) in a geometric space. (**b**) The vector space obtained from core promoters (*V_reguiatory_*) and their corresponding control (*V_random_*) define two different groups of closest *k*-mers (cosine similarity) to the ‘CTATATA’ vector (*v_CTATATA_*). The group of closest *k*-mers in *V_reguiatory_*, when compared to the group formed in *V_random_*, are more similar in sequence (shorter edit distance), and have in average more positive *k*-mer scores from an equivalent “bag-of-*k*-mers” model. This suggests a functional relationship between those *k*-mers in regulatory sequences versus random regions. (**c**) Likewise, the group of *k*-mers closest in the *V_regulatory_* space share positional preferences relative to the TSSs in the region expected for the TATA element. (**d**) Their counterpart in *V_random_* does not.

To illustrate, we compared the representative vector of ‘CTATATA’ in *V_reguiatory_* (*i.e*., set of *v_k-mers_* learned from core promoter regions) and in *V*_random_ (*i.e*., set of *v_k-mers_* learned from random regions used as controls for core promoters). Using *v_CTATATA_* we obtained the set of top five closest *v_k-mers_* in *V_regulatory_* and in *V_random_* and found that *k*-mers from *V_reguiatory_* share more sequence similarity (average edit distance1.8 vs 4.2 respectively) and have, on average, more positive scores from the respective “bag-of-*k*-mers” model (1.49 vs 0.01) (Fig. 6b). In addition, *k*-mers close to *v_CTATATA_* in *V*_regulatory_ share positional constraints that are not recovered from those related in *V_random_* (Fig. 6c-d). This example shows how the output of the geometric spaces can be exploited to determine groups of similar *k*-mers according to their context.

To obtain a global view of how many *k*-mers are embedded in different local sequences between regulatory and random regions, we collected for any given *k*-mer (*k*=8) in the vocabulary, the list of the closest similar *k*-mers ranked by cosine similarity from *V*_regulatory_ and *V*_random_. Next, we contrasted the two ranked lists and determined which *k*-mers show the greatest dissimilarity between regulatory and random regions [41]. In general, we found that low complexity *k*-mers do not show distinctive organizational ‘rules’ between regulatory regions and random, reinforcing our view that short repetitive sequences are not important to define the identity of a sequence. We found that, in terms of the number of *k*-mers with different relationships between *V_regulatory_* and *V_random_*, “vector-*k*-mers” models derived from TF binding loci (~45%) and core promoter regions (~30%) result in notably more differentially represented *k*-mers than models derived from open chromatin regions (~5%) (Supplementary Table 6). In all the cases, we observed a similar proportion of *k*-mers enriched and depleted in regulatory regions (as established from the “bag-of-*k*-mers” scores). The results from models trained in open chromatin regions, might represent the heterogeneity of the regions that prevents the model from learning many specific *k*-mer vectors. However, the fact that the classifiers work with great accuracy indicates that even when the differences are less pronounced than for TF binding loci and core promoter regions, they are large enough to distinguish between an open chromatin region and its control.

We integrated the information obtained from the “bag-of-*k*-mers” and the “vector-*k*-mers” models and found that for the top 1% of the *k*-mers that are enriched in frequency in regulatory regions there is little overlap between *k*-mers that resemble motifs and *k*-mers that show differential relationships between regulatory regions and random regions (Supplementary Table 6). For instance, from the FEA4 models, only 10 out of 103 *k*-mers, that are statistically similar to Arabidopsis motifs, show differential *k*-mer relationships between regulatory and random regions. The difference might be derived from the proportion of TFBSs that are not similar between Maize and Arabidopsis cistromes. In summary, we have compiled a regulatory vocabulary that includes a proportion of key *k*-mers that are enriched in regulatory regions and (1) resemble known motifs, and (2) are embedded in a specific regulatory context.

## DISCUSSION

The decreased cost of large scale genotyping and genome assemblies for crops such as maize and related species, has already shown potential to accelerate the breeding process by linking sequence and structural variation to phenotype [42]. A vast amount of functional genetic variation that is important to phenotype is located in the non-coding regions of the genome. This variation is largely untapped because recognizing functional alleles in the non-coding regions of the genome is both expensive and laborious. In humans and other metazoan models, non-coding annotation that allows identification of functional genetic variation has been accelerated over the last decade using two types of analyses: (1) functional analysis from large collections of biochemical assays; and (2) comparative sequence analysis between reference genomes of closely related species [43]. Yet, in maize, these two types of analyses are particularly challenging. Large collections of biochemical assays remain prohibitive at the scale necessary to cover maize diversity, which is ~20 times more than the diversity found in humans [44]. In addition, comparative sequence analysis requires genome alignment between closely related species, which for maize and its relatives is complicated by the presence of a large number of repetitive sequences in the genome.

In this study, we introduce a computational framework consisting of two type of machine learning models that can accurately classify regulatory regions obtained from functional genomic experiments and random genomic regions. These approaches were borrowed from the fields of natural language processing and information retrieval, and were explicitly chosen to overcome the challenges of annotating intergenic regions in maize. To address highly repetitive sequences and the role of low-complexity regions in maize non-coding regions the “bag-of-*k*-mers” model relies on first filtering out *k*-mers with low-complexity, and next using a sublinear function to transform raw *k*-mer frequencies to down weight *k*-mers that are too frequently observed in a group of sequences and in consequence have less power to discriminate between regulatory and non-regulatory regions. In parallel, the “vector-*k*-mers” model learns local *k*-mer organization from *k*-mer co-occurrence frequencies, which in practice results in a geometric space that allows alignment-free comparisons between sequences [45]. The simultaneous use of two different approaches adds robustness to the predicted annotations, allowing researchers to contrast or combined the results of the two types of models.

Because both models are amenable to interpretation, examination of the learned features offers novel insights about key sequence characteristics that can help to build mechanistic hypotheses to be tested at molecular level, and allow comparison of regulatory programs under the same framework. For instance, both types of models suggest that low complexity *k*-mers are not important for regulatory regions in maize. Also, through modeling MNAseq data we found that open chromatin regions in maize are characteristically organized within poly(dA:dT) tracts flanking G+C rich *k*-mers resembling motifs (Fig. 5a-b). Likewise, from modeling maize KN1 ChIP-seq data and further annotation of regions bound by OSH1, we determined conservation only at the center of binding loci (Fig. 4f). Taken together, our framework can be used beyond the transference of regional annotations, as can easily be extended to *in silico* evaluate the putative effect of sequence variation (*i.e*., SNPs, single nucleotide polymorphisms) in regulatory function from the differences in *k*-mer scores and regulatory probabilities for small groups of *k*-mers.

This work opens many avenues for improving models by adding relevant layers of information. Possible layers to add include: predictions of the 3D structure of regulatory regions, joint modeling of functional genomic data spanning the range of maize diversity to identify general patterns for relevant phenotypes, or even extended across species to build generalizable models that capture conserved features unseen with alignments. Furthermore, we expect these annotations to be useful as priors to improve marker assisted technologies such as genomic selection and to identify targets for genome editing to purge sequence variation contributing to gene expression dysregulation.

## METHODS

### Definition of maize regulatory regions

In the analyses presented throughout this study, we used data sets derived from different functional genomic experiments and obtained from the reference genome (ZmB73 AGPv3, chromosomes 1 to 10) [46]. We included in the analysis open chromatin regions in shoot and roots derived from MNAseq data [3]; binding loci for Knotted 1 (KN1) and Fasciated ear4 (FEA4) transcription factors from ChIP-seq data [27,28], and promoter regions [29–31] from the intersection of TSSs obtained with CAGE and FLcDNAs (Supplementary Table 1). For MNAseq and ChIP-seq peaks, we collected sequences of 300 bp length symmetrically surrounding the midpoints from the originally defined regions. Similarly, for core promoters, we selected the region between −50 bp; +250 bp surrounding the TSSs. Each group of regulatory regions was randomly divided between training and holdout sets and reserved for further analyses.

To randomly select control regions, we first divided the reference genome into sliding windows of 300 bp length, with 50 bp overlap using bedtools v2.24.0 [47] (bedtools makewindows −g zmb73Genome.tsv −w 300 −s 50 > regulatoryRegions.bed) and after removal of all the windows overlapping with the regulatory regions (bedtools −v −a regulatoryRegions.bed −b zmb73GenomeWindows.bed) stored in a sqlite3 database. Next, for each sequence in the training sets we queried the database for regions in their vicinity (10 kb window) with a matching G+C content; if no match was found, we removed the vicinity criteria and searched for a G+C matching region in the same chromosome; if yet no control region could be identified we discarded the regulatory sequence from the training set. For the holdout sets we build balanced and unbalanced holdout sets from randomly selecting one and ten control regions respectively.

### Definition of grasses regulatory regions

Sorghum (*Sorghum bicolor*) core promoter regions were obtained from the reference genome (v2.1) [48] for the coordinates between −50; +250 bp surrounding the start position of genes with annotated 5’UTR and a subset of 1000 sequences randomly selected for further analyses. Rice Knotted1-like (*i.e*., OSH1) binding regions were obtained from re-analyzing ChIP-seq experiment starting with the download of raw data from DDBJ (accession numbers DRA000206 and DR000313) corresponding to two biological replicates of immunoprecipitation with α-OSH1 and IgG antibodies [34]. Raw reads were mapped against the rice reference genome (*Oryza sativa* Nipponbare, IRGSP-1.0 [49] using bowtie v1.1.2 (options −n 2, −l 60, −X 500, --best, --strata, −m 1) [50] and low quality and duplicated reads were removed using picard^1^ (MarkDuplicates) and samtools (−F 780, −F 1024, −f 2) [51] MACS v2.1.0 [52] was used for peak calling (−g 3.73e8, −q 0.01) for each of the replicates and 42 peaks with a reproducible summit reserved and further extended to 300 bp for downstream analyses.

Corresponding control regions were obtained as explained above for maize. Briefly, each reference genome was divided into windows and after removal of sequences overlapping the putative regulatory regions we randomly selected sequences matching G+C content and when possible in the vicinity (~10 kb) of each of the regulatory sequences.

### Preprocessing of sequences

Sequences were preprocessed before fitting models. The preprocessing for the “bag-of-*k*-mers” model involves the dividing of each sequence into 1bp sliding (overlapping) windows of a given size *k* (*k*-mers) to collect for a sequence of length L (L-k)+1 *k*-mers. Next, *k*-mers were converted into tokens (*t*) that correspond to collapsed pairs of *k*-mer and their respective reversed complementary. For the “vector-*k*-mers” models, each sequence is described as a collection of “sentences” resulting from walking *k* times and sliding by 1bp. Each sentence is broken into ordered non-overlapping *k*-mers and next converted into new tokens, as described for the “bag-of-*k*-mers”.

### Calculation of TF-IDF and implementation of the “bag-of-*k*-mers” model

Let’s define all the sequences in a given set from a functional genomics experiment and its corresponding control regions as a collection *S*=*{s_1_, s_2_*,…*s_n_}* of individual sequence *s_i_*=*{t_1_, t_2_*,…, *t_n_}* divided into tokens *t*. The set of all the possible tokens for a given *k* belong to the vocabulary, *Y*. Each *s_i_* is mapped to a list of token weights −*W*_s_- of size |*Y*| that contains “weights” for each token that occurs in *s_i_*, where the “weight” (**equation 1**) is defined as the product of the token frequency - *f*(*t*) - in *s_i_*, and its inverse collection frequency - *idf*(*t*)-. Calculation of TF-IDF were done according to the implementation in the python library scikit-learn v0.19.0 [53].

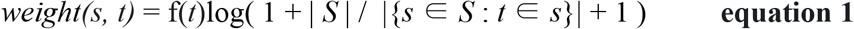

To generate a “bag-of-*k*-mers” model, each training data set is represented as a matrix with *W*_s_-list of token weights- as rows, and a list of sequence labels (1 for regulatory regions and 0 for control regions) to fit a regularized logistic regression were the *C* parameter has been chosen by fivefold cross-validation using a grid search function logistic regression and grid search function used correspond to the implementation of the python library scikit-learn v0.19.0 [53].

### Implementation of “vector-*k*-mers” model

To generate “vector-*k*-mers” models we used the implementation of word2vec algorithms from the python library gensim v1.0.0, which fits sequence representations (*k*-mer vectors - *v*_k-mers_) via Stochastic Gradient Descent (SGD) that aims to optimize an objective function, that implicitly correspond to likelihood for *k*-mer occurrences [54,25]. Next, as shown for text classification, sequence representations −*v*_k-mers_- can be turned through inversion via Bayes rule to determine the likelihood of a new sequence of being part of a regulatory region based on its *k*-mer composition [26]. This classification schema interprets the individual *v*_k-mers_ as components in a composite likelihood approximation that allows classification of sequences without extra modeling or estimation steps.

In brief, we trained a shallow (one single hidden layer), fully connected neural network aimed to optimize the probability of predicting a given *k*-mer (*k*-mertarget) from its context, that is from the observation of the co-occurring *k*-mers appearing anywhere within a small window around the target. We ran word2vec with 30 iterations using hierarchical softmax and no negative sampling (iter=30, hs=1, negative=0, size=300, min_count = 0 and window = 5, all others parameters were kept as the defaults) for each data set and obtain two independent geometric spaces (a continuous space of sequence representations), one for the regulatory regions (*V_reguiatory_*) and the other for the control regions (*V_random_*).

For the classification step, we calculated the probability of every new sequence *s*_i_ under each sequence representation – *V_reguiatory_* and *V_random_* – by first calculating the likelihood of every window within a sentence (using the score function from gensim) and the averaging likelihoods to obtain sentence likelihoods. Next, from the matrix of sentence likelihoods by the two categories (*i.e., C* = regulatory and control) we derive the sequence probabilities - *pV*_regulatory_(*s_i_*) and *pV*_random_(*s_i_*). The category probabilities were calculated via Bayes rule, using as priors *π_C_*=1/C, such that the classification proceeds by assigning the category for which *pV*_category_(*s_i_*) is greater [26].

### Evaluation of models performance

Accuracy, ROC and precision recall curves were generated using the python library scikit-learn v0.19.0 [53] and plotted with python matplotlib v2.0.0 [55].

### Calculation of *k*-mer complexity on a TF motifs database

The sequence complexity of any *k*-mer was approximated to the Shannon entropy for the symbols succession given by equation 2.

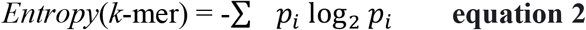

Were *p_i_* correspond to the probability of appearance of the *i*-th symbol in the *k*-mer.

To empirically establish a threshold of complexity for *k*-mers within regulatory regions we calculated the *k*-mer complexity for any given *k* and for all the consensus sequences derived from transcription factor (TF) binding models represented as Position Weight Matrices (PWMs) in the HOmo sapiens COmprehensive MOdel COllection (HOCOMOCO) v10 [56].

### Motif enrichment analyses

To identify *k*-mers similarity to transcription factor binding sites we used TOMTOM from the MEME suite [37] and two collections of *Arabidopsis thaliana* TF binding motifs derived from large-scale experiments [38,39]. The fold enrichment was calculated according to equation 3, in which **N** correspond to the size of the *k*-mer vocabulary, ***n*** correspond to the 1% of the *k*-mer vocabulary taking from the top after sorted with the weights obtained from the model, ***M*** correspond to the number of *k*-mers with a significant hit against a TF motif and ***m*** to the number of *k*-mers that are in the top 1% and have a significant hit against a TF motif.

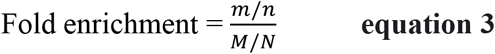

The statistical significance of the enrichment was calculated using the hypergeometric test, as implemented with the python library scipy 0.18.1 (stats.hypergeom), after applying the Bonferroni correction for multiple testing hypothesis to the alpha value required for statistical significance.

## Data accessibility

All the regulatory and control sequences together with the code used to train and evaluate the models are available through Bitbucket repository

## Acknowledgments

This work has been funded by NSF Plant Genome Project (IOS #1238014) USDA-ARS

## Footnotes

1 http://broadinstitute.github.io/picard/

## REFERENCES

1. Wallace JG, Bradbury PJ, Zhang N, Gibon Y, Stitt M, Buckler ES. Association Mapping across Numerous Traits Reveals Patterns of Functional Variation in Maize. Borevitz JO, editor. PLoS Genet; 2014;10:e1004845.

2. Liu H, Luo X, Niu L, Xiao Y, Chen L, Liu J, et al. Distant eQTLs and Non-coding Sequences Play Critical Roles in Regulating Gene Expression and Quantitative Trait Variation in Maize. Molecular Plant. 2017;10:414–26.

3. Rodgers-Melnick E, Vera DL, Bass HW, Buckler ES. Open chromatin reveals the functional maize genome. Proc. Natl. Acad. Sci. U.S.A. 2016;113:E3177–84.

4. Lu F, Romay MC, Glaubitz JC, Bradbury PJ, Elshire RJ, Wang T, et al. High-resolution genetic mapping of maize pan-genome sequence anchors. Nat Commun. 2015;6:6914.

5. Ajmone-Marsan P, Stella A. Commentary on the 6th International Symposium of Animal Functional Genomics. Genet. Sel. Evol. 2016;48:97.

6. Poland J. Breeding-assisted genomics. Curr. Opin. Plant Biol. 2015;24:119–24.

7. Natarajan A, Yardimci GG, Sheffield NC, Crawford GE, Ohler U. Predicting cell-type-specific gene expression from regions of open chromatin. Genome Res. 2012;22:1711–22.

8. Huminiecki L, Horbanczuk J. Can We Predict Gene Expression by Understanding Proximal Promoter Architecture? Trends Biotechnol. 2017;35:530–546.

9. Stringham JL, Brown AS, Drewell RA, Dresch JM. Flanking sequence context-dependent transcription factor binding in early Drosophila development. BMC Bioinformatics. 2013;14:298.

10. Stampfel G, Kazmar T, Frank O, Wienerroither S, Reiter F, Stark A. Transcriptional regulators form diverse groups with context-dependent regulatory functions. Nature. 2015;528:1470–51

11. Crocker J, Abe N, Rinaldi L, McGregor AP, Frankel N, Wang S, et al. Low Affinity Binding Site Clusters Confer Hox Specificity and Regulatory Robustness. Cell. 2015;160:191–203.

12. Raveh-Sadka T, Levo M, Shabi U, Shany B, Keren L, Lotan-Pompan M, et al. Manipulating nucleosome disfavoring sequences allows fine-tune regulation of gene expression in yeast. Nature Genetics. 2012;44:743–50.

13. Farley EK, Olson KM, Zhang W, Rokhsar DS, Levine MS. Syntax compensates for poor binding sites to encode tissue specificity of developmental enhancers. Proc. Natl. Acad. Sci. U.S.A. 2016;113:6508–13.

14. Yáñez-Cuna JO, Kvon EZ, Stark A. Deciphering the transcriptional cis-regulatory code. Trends Genet. 2013;29:11–22.

15. Lee D, Karchin R, Beer MA. Discriminative prediction of mammalian enhancers from DNA sequence. Genome Res. 2011;21:2167–80.

16. Lee D, Gorkin DU, Baker M, Strober BJ, Asoni AL, McCallion AS, et al. A method to predict the impact of regulatory variants from DNA sequence. Nature Genetics. 2015;47:955–61.

17. Ghandi M, Lee D, Mohammad-Noori M, Beer MA. Enhanced Regulatory Sequence Prediction Using Gapped k-mer Features. Morris Q, editor. PLoS Comput Biol. 2014;10:e1003711.

18. Alipanahi B, Delong A, Weirauch MT, Frey BJ. Predicting the sequence specificities of DNA-and RNA-binding proteins by deep learning. Nature Biotechnology. 2015;33:831–8.

19. Zhou J, Troyanskaya OG. Predicting effects of noncoding variants with deep learning-based sequence model. Nat. Methods. 2015;12:931–4.

20. Kelley DR, Snoek J, Rinn JL. Basset: learning the regulatory code of the accessible genome with deep convolutional neural networks. Genome Res. 2016;26:990–9.

21. Zhang D, Wang D. Relation Classification: CNN or RNN? Natural Language Understanding and Intelligent Applications. 2016. pp. 665–75.

22. Yin W., Kann, K., Yu, M., & Schutze. Comparative Study of CNN and RNN for Natural Language Processing. arXiv. 2017;1702.01923

23. Manning CD, Schütze H. Foundations of Statistical Natural Language Processing. MIT Press; 1999.

24. Mikolov T, Sutskever I, Chen K, Corrado GS, Dean J. Distributed Representations of Words and Phrases and their Compositionality. arXiv. 2013;1310.4546.

25. Mikolov T, Chen K, Corrado G, Dean J. Efficient Estimation of Word Representations in Vector Space. arXiv. 2013; 1301.3781.

26. Taddy M. Document Classification by Inversion of Distributed Language Representations. arXiv. 2015; 1504.07295.

27. Bolduc N, Yilmaz A, Mejia-Guerra MK, Morohashi K, O’Connor D, Grotewold E, et al. Unraveling the KNOTTED1 regulatory network in maize meristems. Genes Dev. 2012;26:1685–90.

28. Pautler M, Eveland AL, LaRue T, Yang F, Weeks R, Lunde C, et al. FASCIATED EAR4 encodes a bZIP transcription factor that regulates shoot meristem size in maize. The Plant Cell. 2015;27:104–20.

29. Alexandrov NN, Brover VV, Freidin S, Troukhan ME, Tatarinova TV, Zhang H, et al. Insights into corn genes derived from large-scale cDNA sequencing. Plant Mol. Biol. 2009;69:179–94.

30. Soderlund C, Descour A, Kudrna D, Bomhoff M, Boyd L, Currie J, et al. Sequencing, mapping, and analysis of 27,455 maize full-length cDNAs. PLoS Genet. 2009;5:e1000740.

31. Mejia-Guerra MK, Li W, Galeano NF, Vidal M, Gray J, Doseff AI, Grotewold E. Core Promoter Plasticity Between Maize Tissues and Genotypes Contrasts with Predominance of Sharp Transcription Initiation Sites. The Plant Cell. 2016;27:3309–20.

32. Liu Q, Gan M, Jiang R. A sequence-based method to predict the impact of regulatory variants using random forest. BMC Syst Biol. 2017; 11:7.

33. Bolduc N, Hake S. The maize transcription factor KNOTTED1 directly regulates the gibberellin catabolism gene ga2ox1. The Plant Cell. 2009;21:1647–58.

34. Tsuda K, Kurata N, Ohyanagi H, Hake S. Genome-wide study of KNOX regulatory network reveals brassinosteroid catabolic genes important for shoot meristem function in rice. The Plant Cell. 2014;26:3488–500.

35. Wang J, Zhuang J, Iyer S, Lin X, Whitfield TW, Greven MC, et al. Sequence features and chromatin structure around the genomic regions bound by 119 human transcription factors. Genome Res. 2012;22:1798–812.

36. Dror I, Rohs R, Mandel-Gutfreund Y. How motif environment influences transcription factor search dynamics: Finding a needle in a haystack. Bioessays. 2016;38:605–12.

37. Gupta S, Stamatoyannopoulos JA, Bailey TL, Noble WS. Quantifying similarity between motifs. Genome Biol. 2007;8:R24.

38. Franco-Zorrilla JM, López-Vidriero I, Carrasco JL, Godoy M, Vera P, Solano R. DNA-binding specificities of plant transcription factors and their potential to define target genes. Proc. Natl. Acad. Sci. U.S.A. 2014;111:2367–72.

39. O’Malley RC, Huang S-SC, Song L, Lewsey MG, Bartlett A, Nery JR, et al. Cistrome and Epicistrome Features Shape the Regulatory DNA Landscape. Cell. 2016;165:1280–92.

40. Levy O, Goldberg Y. Linguistic Regularities in Sparse and Explicit Word Representations. Proceedings of the Eighteenth Conference on Computational Natural Language Learning. 2014;171–80.

41. Webber W, Moffat A, Zobel J. A similarity measure for indefinite rankings. ACM Transactions on Information Systems. 2010;28:1–38.

42. Jiao Y, Peluso P, Shi J, Liang T, Stitzer MC, Wang B, et al. Improved maize reference genome with single-molecule technologies. Nature. 2017;546:524–7.

43. Alexander RP, Fang G, Rozowsky J, Snyder M, Gerstein MB. Annotating non-coding regions of the genome. Nature Reviews Genetics. 2010;11:559–71.

44. Buckler ES, Gaut BS, McMullen MD. Molecular and functional diversity of maize. Curr. Opin. Plant Biol. 2006;9:172–6.

45. Asgari E, Mofrad MRK. Continuous Distributed Representation of Biological Sequences for Deep Proteomics and Genomics. PLoS ONE. 2015;10:e0141287.

46. Schnable PS, Ware D, Fulton RS, Stein JC, Wei F, Pasternak S, et al. The B73 maize genome: complexity, diversity, and dynamics. Science. 2009;326:1112–5.

47. Quinlan AR. BEDTools: The Swiss-Army Tool for Genome Feature Analysis. Curr Protoc Bioinformatics. 2014;47:11.12.1-34.

48. Paterson AH, Bowers JE, Bruggmann R, Dubchak I, Grimwood J, Gundlach H, et al. The Sorghum bicolor genome and the diversification of grasses. Nature. 2009;457:551–6.

49. Kawahara Y, la Bastide de M, Hamilton JP, Kanamori H, McCombie WR, Ouyang S, et al. Improvement of the Oryza sativa Nipponbare reference genome using next generation sequence and optical map data. Rice (N Y). 2013;6:4.

50. Langmead B, Trapnell C, Pop M, Salzberg SL. Ultrafast and memory-efficient alignment of short DNA sequences to the human genome. Genome Biol. 2009;10:R25.

51. Li H, Handsaker B, Wysoker A, Fennell T, Ruan J, Homer N, et al. The Sequence Alignment/Map format and SAMtools. Bioinformatics. 2009;25:2078–9.

52. Zhang Y, Liu T, Meyer CA, Eeckhoute J, Johnson DS, Bernstein BE, et al. Model-based analysis of ChIP-Seq (MACS). Genome Biol. 2008;9:R137.

53. Pedregosa, F., Varoquaux, G., Gramfort, A.., Michel, V. Thirion, B., Grisel, O.., Blondel, M., Prettenhofer, P., Weiss, R., Dubourg, V., Vanderplas, J., Passos, A., Cournapeau, D., Brucher, M., Perrot, M., Duchesnay, E. Journal of Machine Learning Research. 2011;12:2825–2830.

54. Rehurek, R., & Sojka, P. Software Framework for Topic Modelling with Large Corpora. in Proceedings of the LREC 2010 Workshop of New Challenges for NLP Frameworks. 2010; 45–50

55. Hunter JD. Matplotlib: A 2D Graphics Environment. Computing in Science & Engineering. 2007;9:90–5.

56. Kulakovskiy IV, Vorontsov IE, Yevshin IS, Soboleva AV, Kasianov AS, Ashoor H, et al. HOCOMOCO: expansion and enhancement of the collection of transcription factor binding sites models. Nucleic Acids Res. 2016;44:D116–25.

